# DCATS: differential composition analysis for complex single-cell experimental designs

**DOI:** 10.1101/2022.03.21.485232

**Authors:** Xinyi Lin, Chuen Chau, Yuanhua Huang, Joshua W.K. Ho

## Abstract

Differential composition analysis – the identification of cell types that have statistically significantly change in abundance between multiple experimental conditions – is one of the most common tasks in single cell omic data analysis. However, it remains challenging to perform differential composition analysis in the presence of complex experimental designs and uncertainty in cell type assignment. Here, we introduce a statistical model and an open source R package, DCATS, for differential composition analysis based on a beta-binomial regression framework that addresses these challenges. Our empirical evaluation shows that DCATS consistently maintain high sensitively and specificity compared to state-of-the-art methods.

## 1 Introduction

Single-cell RNA sequencing (scRNA-seq) is a high-throughput sequencing technology that probes transcriptomes of a large number of individual cells, allowing researchers to characterise different cell types in a heterogeneous population. It plays a critical role in strengthening our understanding of various biological systems including embryogenesis, the development of different diseases and how cells react to environmental stimuli [1, 2]. Recently, highly multiplexed strategies have been introduced to mix samples from different donors, conditions or treatments with external molecular barcodes or intrinsic genetic makeups to achieve higher efficiency and lower batch effects [3, 4, 5]. In such multi-sample designs, analysis of the differential composition of cell types between two conditions is routinely applied.

From a scRNA-seq experiment, we can obtain the cell counts ***N*** = {*n_1_*,…,*n_K_*}of *K* cell types by performing cell clustering, e.g., with Louvain (a graph-based method) [6] or K-means, usually on reduced dimensions through principal component analysis. Then, the obtained cell count vector ***N*** is conventionally used to estimate the cell type composition abundance ***μ***, with the assumption that the clustering is unbiased and accurate. Thus, a few statistical methods can be directly applied for detecting the cell types with differential composition abundance between conditions, e.g., Fisher exact test. Some statistical tools are also developed specialized for scRNA-seq data. scCODA [7] assumes cell counts of different cell types follows hierarchical Dirichlet-Multinomial distribution. This allows scCODA to model all cell type together. MiloR [8] evaluates differential abundance on smaller clusters in KNN graph which are called ‘neighborhoods’. It assumes that cell counts follow a negative binomial distribution and uses representative cells instead of all cells to improve program efficiency. DAseq [9] also makes use of KNN graph. It calculates multiscale differential abundance score by counting numbers of cells coming from different biological states while varying k. This multiscale differential abundance score is what DAseq used to infer cell states that have differential abundance.

However, it remains a statistical challenge for multiple reasons. First, multi-level variability exists due to technical and biological reasons, for example, low number of biological replicates or low number of cells for minor cell types. Fisher exact test usually can handle the variability sourced from low cell counts through a binomial noise model, but it cannot fully utilise the biological replicates (if any) to account for biological variability. Recently, scDC [10], a Poisson regression-based method, has been introduced to account for the uncertainty not only between replicates but also clustering by leveraging a bootstrap re-sampling, preferred with re-clustering. However, the Poisson noise model is not able to capture the over dispersion. Second, misclassification during the cell clustering step may introduce both systematic bias and uncertainty. For example, subtypes of T helper cells are often confused between each other. A similar challenge was also noticed in meta-genomics analysis, where species with similar sequences are often confusedly aligned and quantified, and bias correction by reversing this bias was found beneficial for the differential abundance analysis [11, 12]. Regarding to challenges mentioned above, we introduced a statistical method, DCATS, to effectively detect the cell types with differential abundance between conditions or along with continuous covariates. This algorithm is implemented as an R package named DCATS which is available at https://github.com/holab-hku/DCATS.

## 2 Results

### 2.1 High-level description of DCATS

DCATS has unique features in two folds. First, it considers the uncertainty of cell type assignment by the biased similarity between cells types. The latent true cell type proportion is obtained by leveraging a similarity matrix between cell types through a maximum likelihood estimation. Second, we introduced a beta-binomial regression model to perform the differential cell type abundance analysis. This model has the benefits to both model the raw cell counts rather than the normalized proportions and consider the dispersion between samples (Fig. 1A; Methods). The effectiveness of the vanilla beta-binomial regression was demonstrated in [13]. However, the estimation of the over dispersion term in this task is often challenging due to the extremely low numbers of replicates (sometimes even single replicate). Therefore, we introduce a strategy to estimate the dispersion term for all cell types jointly, through a pooled beta-binomial regression in a pre-step (Methods).

**Figure 1:**
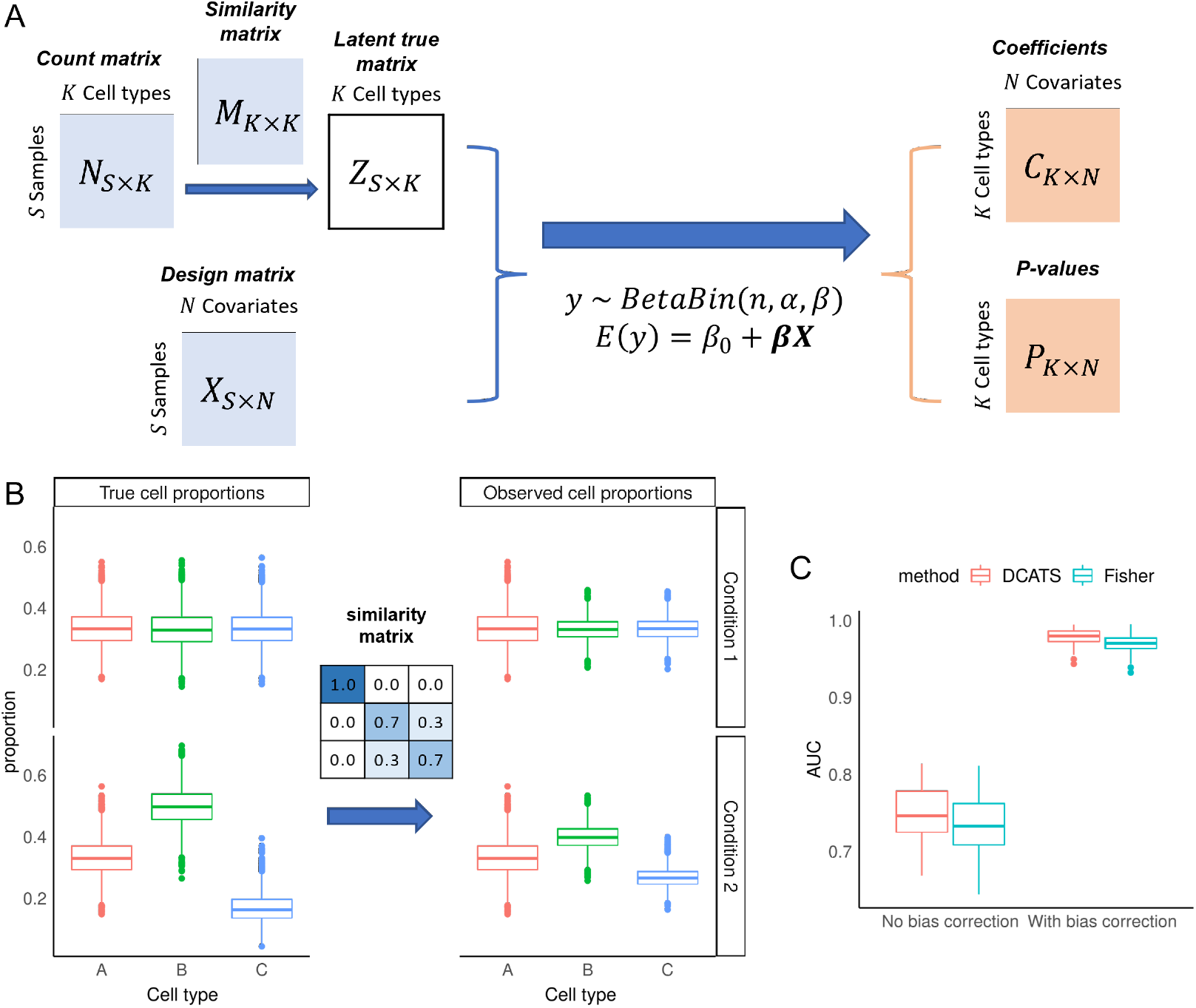
DCATS improves composition analysis through accounting for uncertainty in classification of cell types in differential abundance analysis. A) Illustration of the DCATS workflow. Matrices with light blue are input matrices, matrices with light orange are outputs from DCATS. First step is bias correction using similarity matrix. This step is optional. The design matrix with multiple covariates is also required. DCATS supports both categorical and continuous covariates. Then, DCATS detects differential abundance using a beta-binomial generalized linear model (GLM) model, which returns the estimated coefficients and p-values. B) These box plots illustrate the effect of cell type misclassificaiton in a theoretical simulation with Dirichlet-Multinomial sampling. The similarity matrix is designed to introduce misclassification errors. The proportions of cell types B and C are changed between conditions 1 and 2. C) Area under the precision-recall curve (AUC, same below) values when varying p value in detecting the cell type with differential abundance.

When analyzing scRNA-seq data, we started from gene expression matrix of read or unique molecular identifier (UMI) counts. After basic pre-processing including filtering cells, normalization, and integration [14, 15], several methods can be used to annotate each cells, including manual annotation, supervised methods and semi-supervised methods [16, 17]. The input cell count matrix for DCATS can be calculated by counting the numbers of cells in each cell type.

In order to estimate the cell type similarity matrix *M* for the confusion during clustering, we introduced two heuristic methods here. One is based on the KNN graph between cells by counting the averaged frequency of cell types in each cell’s neighbors (Methods), where the KNN graph may be pre-computed if using Seurat [18] or Scanpy [19] pipelines. The other is to use a prediction confusion matrix produced by a classifier on top principle components, e.g., support vector machine as default.

### 2.2 Correcting misclustering improves composition analysis

To evaluate the effects of misclustering on the composition analysis, we performed a theoretical simulation by generating cell counts from Dirichlet-Multinomial distributions, where the input cell type proportion are generated by adding bias to the genuine cell type proportions through a transformation with the misclustering matrix, aka similarity matrix (Fig. 1C, Supp. Fig. S1).

Here, the genuine proportions of the three cells types are [1/3, 1/3, 1/3] and [1/3, 1/2, 1/6] in conditions 1 and 2, respectively. However, due to the high similarity between cell types 2 and 3, the average cell type proportions are transformed to [0.33, 0.33, 0.33] and [0.33, 0.40, 0.27] respectively in the two conditions, which is mimicking the bias introduced in the clustering. Due to the equal cell type proportion and symmetrical similarity matrix, all cell type proportions remain unchanged in condition 1 after the similarity-based transformation. However, the proportion of cell type 2 and cell type 3 in condition 2 changed due to the mis-classification. Particularly, the difference of proportions between conditions are reduced for both cell types 2 and 3, which may cause false negatives due to the shrunken effects. Indeed, both Fisher exact test and DCATS only return moderate performance (AUC = 0.721 and 0.745 in average, aggregating 50 runs). By contrast, we found that the performance in detecting differential abundance are dramatically increased for both DCATS (mean AUC: 0.745 to 0.976) and Fisher exact test (mean AUC: 0.721 to 0.961; Fig. 1C) by using the cell counts corrected from similarity matrix by DCATS through an Expectation-Maximization algorithm. This is largely thanks to the improved sensitivity in both methods (Supp. Fig. S2).

### 2.3 Benchmarking DCATS with simulated data

We further benchmarked DCATS’s performance with five existing methods: Fisher exact test, scDC [10], speckle [20], diffcyt (a method for similarity purpose but on mass cytometry) [21], and milo [8]. Here, we first generated a large pool of single-cell transcriptomic profiles with a simulator splatter [22] (Fig. 2A; Supp. Fig. S3). This simulated pool is then used as the seed data and cells are randomly selected according to the simulated cell-type proportions (as ground truth; see more details in Methods). Clustering is further performed on the subsets of simulated cells with Seurat to mimic the potential confusion introduced in the clustering step (the empirical confusion matrix shown in Supp. Fig. S5). The six methods are then performed on the cell counts obtained from the clustering annotation. Uniquely, we estimated the similarity matrix between cells using KNN graph by calculating the fraction of neighbours for all cells in each cell type that belong to different cell types. It approximates the empirical confusion matrix well (Pearson’s R=0.9987; Mean absolute error=2.977 × 10^−18^; Supp. Fig. S6).

**Figure 2:**
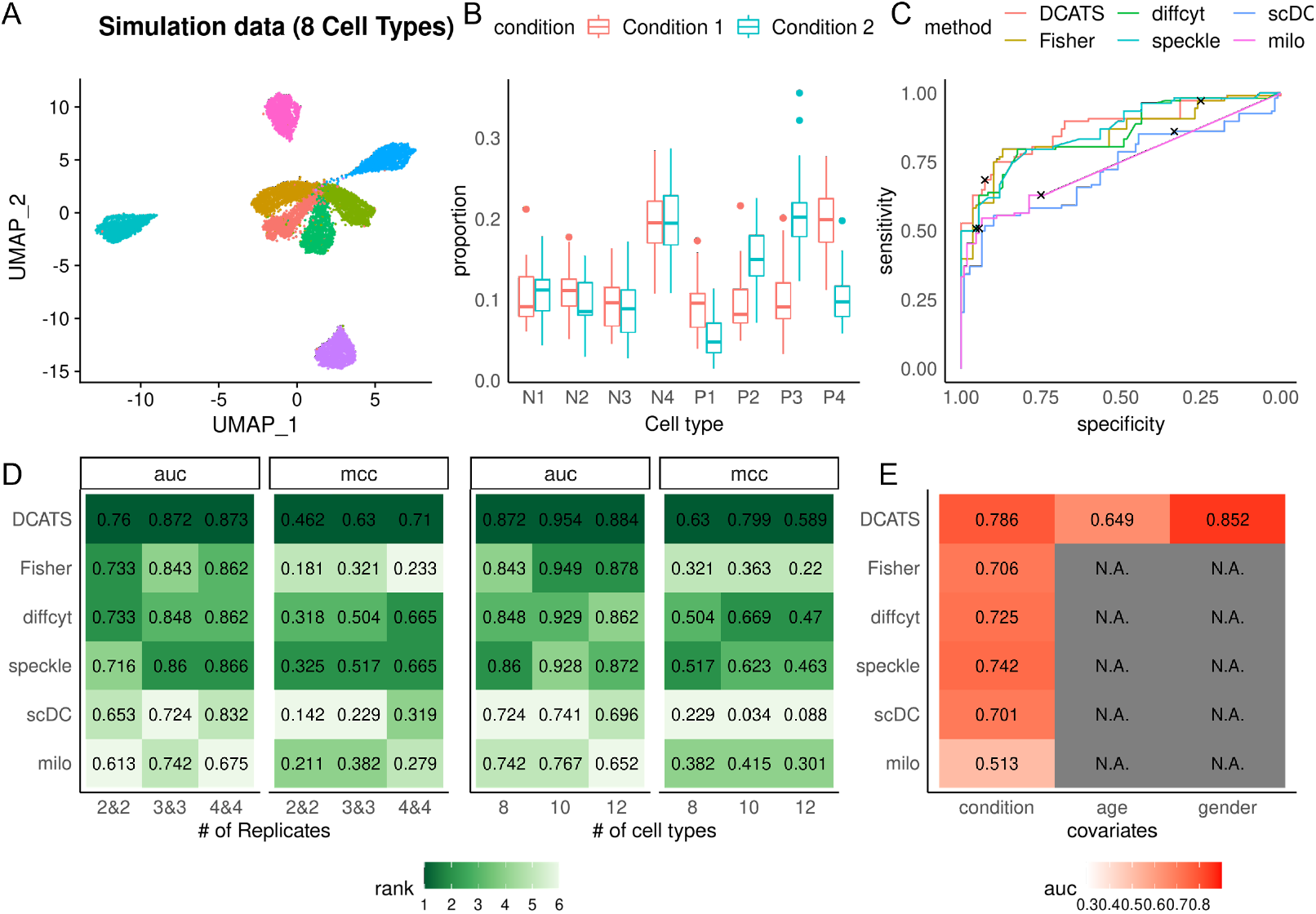
Evaluation of DCATS with multiple simulation datasets. A) The UMAP plots of simulation data with 8 cell types. B) The true proportions of different cell types in default setting (3 replicates in each condition across 30 runs; the 4 cell types with differential composition abundance have names started with ‘P’). C) The ROC curves of different methods in the default setting. Black dots indicating the sensitivities and specificities calculated using defined cutoff (perturbation level = 0 for Milo, p-value = 0.1 for other methods). D) Comparing multiple methods with simulated transcriptomics from Splatter by varying number of replicates (left) and number of cell types (right). The default numbers of cell types is 8, and the default number of replicates is 3. E) Detecting variable cell types with multiple covariates, for both categorical and continuous types. Data is simulated with Splatter. N.B., only DCATS supports multiple covariates.

In a default setting with 3 replicates in each condition and 4 out of 8 cells types with differential composition abundance, we found that DCATS outperforms all other methods (AUC: 0.872 vs 0.86; MCC: 0.63 vs 0.517; F1: 0.779 vs 0.714, respectively for DCATS and the second-best method in each metric; Fig. 2D and Supp. Table S1, Supp. Fig. S4). N.B., DCATS remains the best-performed method in all three metrics while the second-best method varies among the alternative methods. By ablating the correction of misclustering in DCATS and introducing over-dispersion term in the beta-binomial regression, we observed that both bias correction and global estimation of over-dispersion contribute to the improvement of overall performance (Supp. Table S1, Supp. Fig. S4). When varying the number of replicates to 2 or 4, or the number of cell types to 10 or 12, we found that DCATS consistently outperforms all other methods, mostly by a large margin (Fig. 2D, Supp. Table S1-S2). Unsurprisingly, the increase of replicates improves the performance for almost all methods, especially for DCATS, suggesting its capability of estimating biological variability from replicates. It is worth mentioning that despite Milo’s unsatisfactory performance in cell type levels, as a tool design for detecting perturbation of cell states in partially overlapping neighborhood, it shows its strength in differential abundance analysis at neighborhood level compared to DCATS and speckle. (Supp. Fig. S7-8)

As a generalised linear model, DCATS has the capability to account for additional covariates or to test multiple covariates jointly in the association with composition abundance for each cell type. To demonstrate this, we simulated 10 replicates in each condition with different ages and genders. There are 8 differential abundant cells types between conditions, 4 of which also have effects from both age and gender. Among all these methods, DCATS is the only method designed to support additional covariates testing simultaneously with consistent high performance. scDC, as a GLM based method, is in principle able to support covariates but the interface is not implemented. Milo, which is based on a negative binomial GLM model, also fails to control influence of confounding covariates although we included related information when performing the analysis. We found DCATS outperforms all other methods, partly thanks to its capability of jointly modelling additional covariates. Indeed, DCATS shows good performance in detecting cell types with additional covariates for both continuous (age) and discrete (gender) variables (Fig. 2E, Supp. Fig. S9-11).

### 2.4 Evaluating DCATS on experimental data sets

To further illustrate the performance of DCATS, we applied DCATS and four other methods (Fisher, scDC, speckle and milo) on three experimental datasets. As we don’t have the KNN matrices used to annotate cell types in these datasets, similarity matrices are calculated by performing a classification types with SVM over the original cell type annotation. We first assessed their performance on controlling false positives by applying them to a negative control data set on PBMCs, where no cell type was reported to have significant proportion change between 8 lupus patients treated with interferon(IFN)-*β* and 8 control samples [4]. Here, we found that DCATS and speckle both have good control of false positives, with no cell types having *p* < 0.1 (Fig. 3A). Similarly, Milo also has a good control on false positives when manually setting its threshold to 0.2 based on the prior knowledge (this threshold is used throughout all data sets below; Supp. Table S3). On the other hand, Fisher exact test severely suffers from type I errors (4 out of 8 cell types passing a significance level at *p* < 0.01); scDC also returns three cell types with *p* < 0.1, including B cells with p < 0.01.

**Figure 3:**
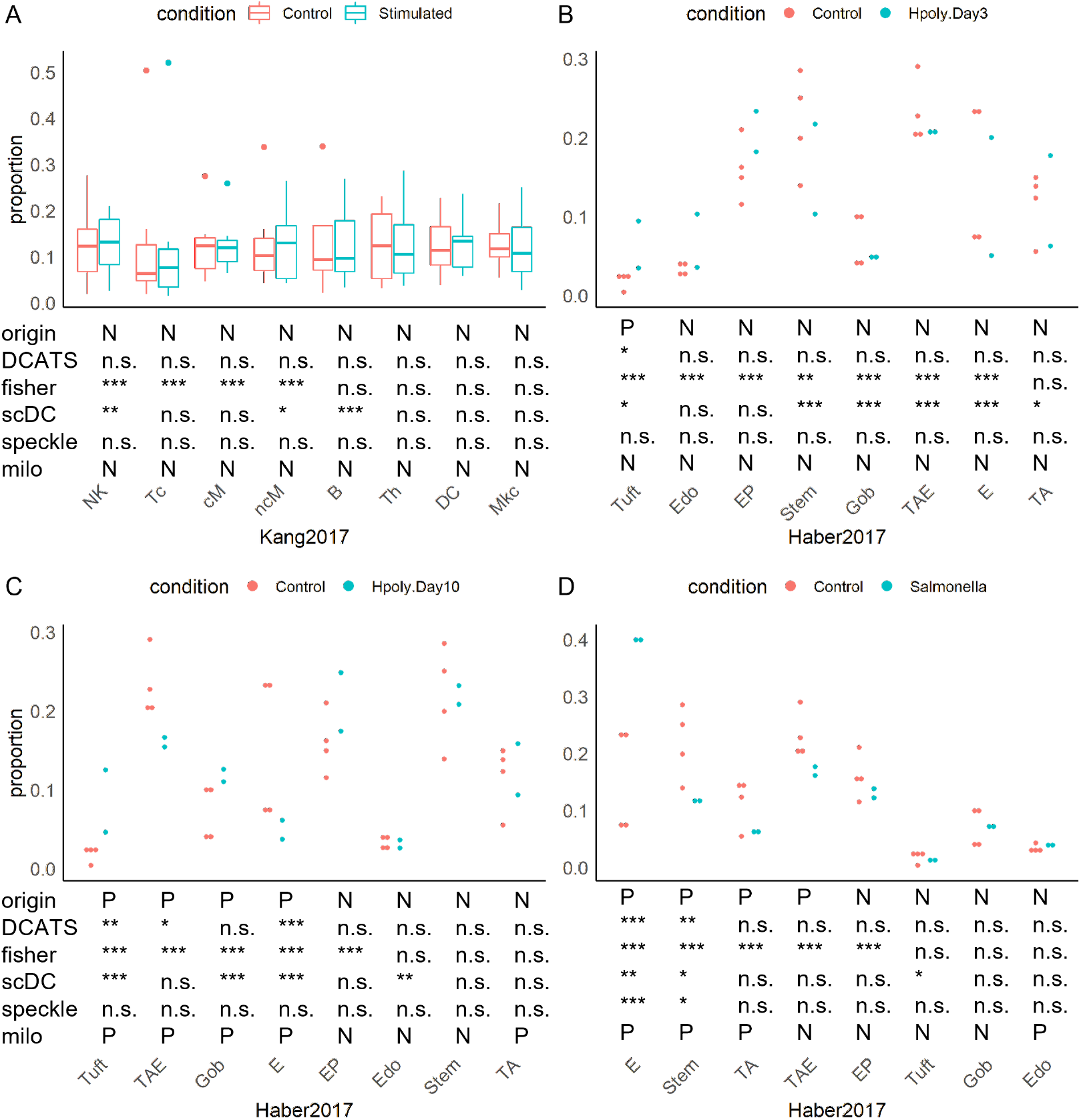
DCATS gives accurate conclusions for two experimental datasets. A) shown is the proportion of each cell types in the Kang dataset [4]. cM, CD14+CD16– monocytes; ncM, CD14+CD16+ monocytes; DC, dendritic cells; Mkc, megakaryocytes; Th, CD4+ T cells; B, B cells; Tc, CD8+ T cells; NK, natural killer cells. B-D) show the proportion of each cell types in the Haber dataset [23]. E, Enterocyte; TA, transit amplifying; TAE, TA.Early; EP, Enterocyte.Progenitor; Gob, Goblet. ‘P’ in the first line represents existing proportions’ difference according to the original papers, and ‘N’ represents no significant proportions’ difference. ‘P’ in the last line represents existing proportions’ difference based on results of milo, and ‘N’ represents no significant proportions’ difference. In the rest four lines, ‘*’ represents p-values from 0.05 to 0.1, ‘**’ represents p-values from 0.01 to 0.05, ‘***’ represents p-values less than 0.01. ‘n.s.’ means not significant

The second dataset consists of 53,193 epithelial cells from mice’s small intestine and organoids [23]. It has 4 control samples, 2 samples coming from two days after *Salmonella* infection, 2 samples coming from three days after *H.polygyrus* infection, and 2 samples coming from ten days after *H.polygyrus* infection. Eight cell types were defined in the original study and then were compared between controls and each stimulation group to identify cell types with differential composition abundance through a Poisson regression and Wald test [23]. Presumably because the sample variability is not well considered in the Poisson regression, the authors used `FDR` < 1e – 5 as the cutoff of differential abundant cell types between control and simulations. When re-analysing this data set, we first noticed that Fisher exact test captured all the reported differential abundant cell types, but also returns false positives in each of the comparison pairs (with *p* < 0.01), particularly between control and Hpoly.Day3 (Fig. 2B-D). Similarly, scDC and milo also returns false positives in each comparing condition pair but occasionally missed reported positives. Surprisingly, speckle shows limited power with missing almost all hits. In contrast, DCATS shows a good balance between sensitivity and specificity; it does not return any false positive (even at a lenient significance level 0.1) and captures most of the reported differential abundant cell types (Fig. 3B-D, Supp. Table S4-5).

### 2.5 Application of DCATS in complex experiment designs

To demonstrate the utility of DCATS for complex designs, we selected a cohort study including 196 hospitalized COVID-19 patients with moderate or severe disease, corresponding control group, and patients in the recovering stage. Within this cohort study, we selected scRNA-seq data from fresh and frozen PBMC samples with known metadata. The samples were stratified into five groups: 20 samples from the control group, 48 samples from the mild or moderate convalescence group, 19 samples from the mild or moderate progression group, 36 samples from the severe or critical convalescence group, and 48 samples from the severe or critical progression group [24]. In the original study, the differential abundance analysis was performed between multiple severity groups.

Different from the original study, we leveraged DCATS’s capability to account for confounding factors, including age and gender through the GLM framework. We noticed both DCATS and the original study identified a few cell types with substantial proportion changes (denoted in black asterisk in Fig. 4), for example, CD8+ T cells between severe in progression and convalescence or healthy controls. On the other hand, DCATS also uniquely detects several cell types that show significant proportion difference (highlighted in red).

**Figure 4:**
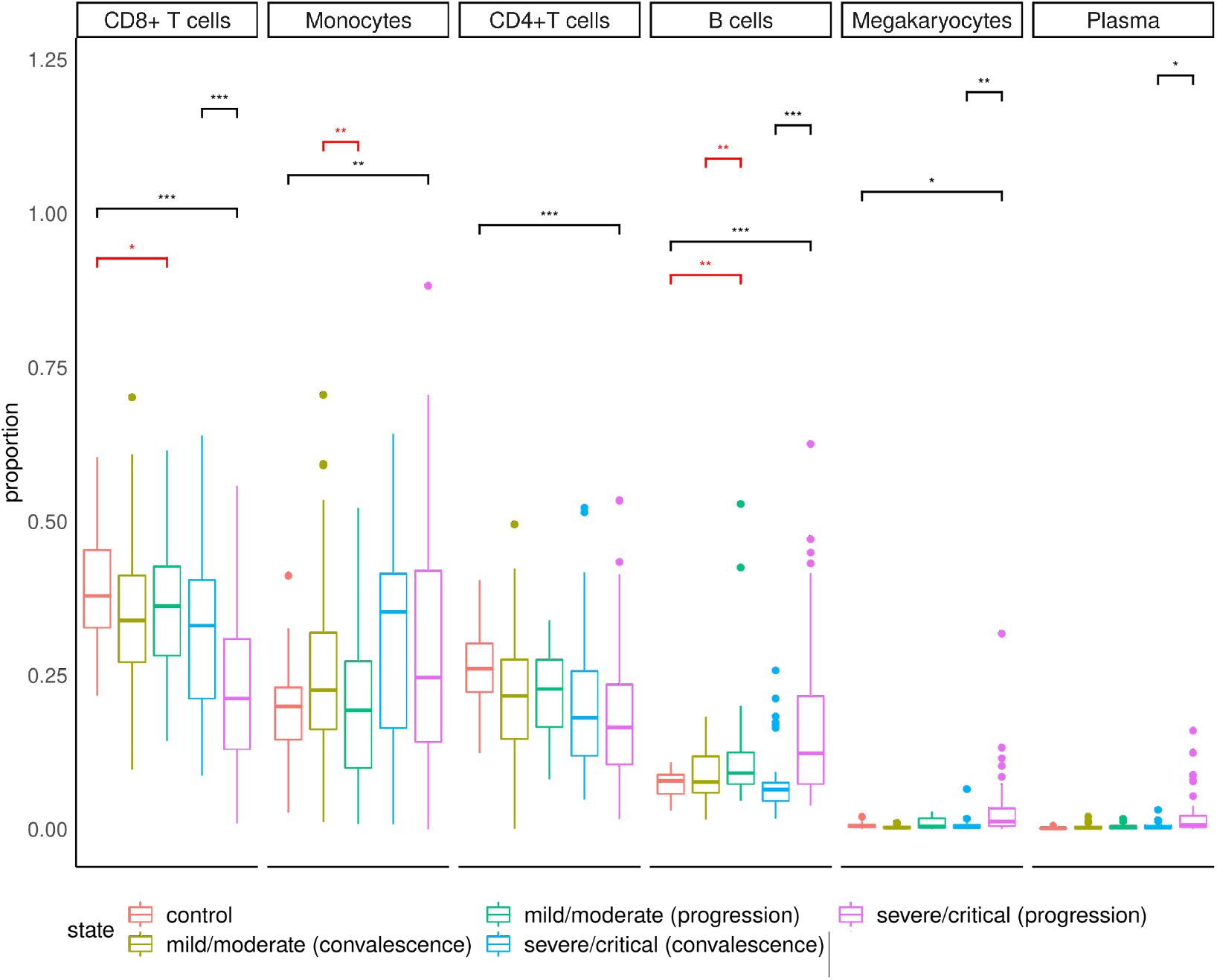
DCATS is able to find new cell types with differential abundance in Ren dataset[24].Cell types’ proportion and conclusions on the Ren dataset given by DCATS. The significant bars in black color indicate that results are the same as the original paper, while the significant bars in red colors indicate newly discovered cell types with proportion difference.

Specifically, DCATS indicates a slightly different proportions of CD8+ T cells between the control group and mild/moderate progression group (*p* = 0.067). This is consistent with what another study found when analyzing high-dimensional cytometry data from 125 COVID-19 patients, corresponding recovered group, and healthy control group [25]. Even though the significance is moderate, the trend is strong. We also observed a significantly difference of proportion in monocytes between mild/moderate progression group and corresponding recovery group (*p* = 0.013). This finding is consistent with a recent report from Qin and colleagues by using fluorescence-activated flow cytometry analysis to track the dynamic changes of monocytes in patients during recovery stage from mild symptoms [26]. DCATS also shows a significant difference in proportion between control group and mild/moderate progression group (*p* = 0.024) as well as the mild/moderate progression group and corre-sponding recovery group (*p* = 0.021) regarding to B cells. The same influence in B cells is also described in the original paper with stronger signal given by DCATS between control group and severe/critical progression group (*p* = 3.10 × 10^-4^) as well as the severe/critical progression group and corresponding recovery group (*p* = 2.48 × 10^-6^). These new findings show that DCATS is powerful tool in identifying cell types with significant proportion changes while keeping a high specificity.

## 3 Discussion and conclusion

In this works, we introduced DCATS, in a form of beta-binomial regression, for detecting cell types with differential composition abundance. With the bias correction procedure based on a similarity matrix, DCATS provides a more robust framework for differential composition analysis between conditions. As the input of DCATS is the count matrix which includes the number of cells in each cell type for each sample, it can deal with pre-defined cell type annotation, and can be easily adapted to any other single cell (multi-) omics data. This unique feature also allows DCATS to be implemented in data with self-adjusted annotation based on integration of multi-omics data or biological knowledge.

Comparing to simple statistical tests like Fisher’s exact test, DCATS preserves high sensitivity while has a good performance in specificity. Through multiple simulation and experimental datasets, we demonstrated that DCATS has an excellent overall performance and outperforms existing methods. Strikingly, DCATS has a strong control of false positives but remains a high level of power.

When benchmarking with other methods, Milo shows unsatisfactory performance when we conducted the differential analysis at cell type level. This is partly because that Milo is designed at the neighborhood level, and there is no build-in metric to determine the differential abundance at cell type level, hence the proportion of neighborhoods with significant difference were used as a surrogate to indicate cell-type level significance. Despite the unsatisfactory performance for Milo, it shows its strength in detecting differential abundance neighborhoods comparing to other methods, and is a good choice when we have data with continuous cell state changes.

As DCATS corrects the misclassification bias based on the similarity matrix, the estimation of this matrix is an important step and can influence the performance of DCATS. Although, we found the KNN-based or empirical classification based similarity matrices function well in general, the estimation of this matrix may be further improved in future, in the light of more benchmarking datasets with accurate cell type annotations.

## 4 Method

### 4.1 Composition correction with clustering similarity matrix

From a scRNA-seq experiment, we can obtain the cell counts ***x*** = {*x*_1_,…,*x_K_*} of *K* cell types by performing cell clustering, e.g., Louvain method [6] and K-means. If the clustering is unbiased and accurate, it is natural to take the cell type composition abundance ***μ*** proportional to the returned cell count vector ***x***. However, due to the complex underlying structure, the clustering step is often inaccurate and may be systematically biased, hence it is crucial to correct the bias on composition abundance introduced by clustering method. In next subsection, we demonstrate that the clustering similarity matrix *M* (or similarity matrix) can be well approximated, e.g., by KNN graph. Specifically, the element *m_i,j_* denotes the probability of a cell *c* in type *i* assigned to type *j* by the clustering method, namely, *P*(*A_c_* = *j*|*I_c_* = *i*) = *m*_*i,j*_, where *A_c_*, *I_c_* respectively denote the clustering assignment and genuine identity of cell *c*. This also means that *M* could be asymmetric and requires 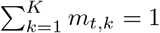 with *m_t,k_* ≥ 0 for any cell type *t*.

Given the above definition and a pre-defined clustering similarity matrix *M*, we could have the likelihood of unknown cell type composition vector ***μ*** on observing the cell counts ***x***, as follows,

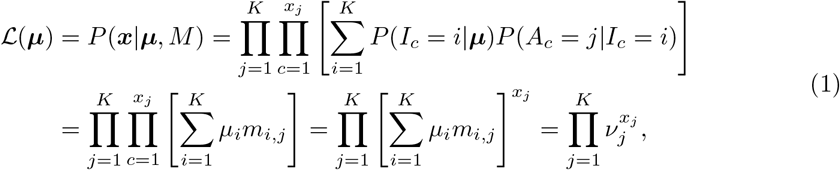

which is equivalent to a multinomial distribution parametrised by an adjusted composition abundance vector ***ν*** = **μ** × *M*.

Algebraically, we could take **μ** = M^-1^ × **ν** with inverting the similarity matrix *M*. However, this simplistic treatment does not guarantee that the solution satisfies with *μ_t_* ≥ 0 and *μ_t_* ≤ 1 for any cell type *t*. In practice, constraints have been introduced to address this optimisation problem, e.g., [12, 11], though it occasionally returns unstable solutions.

Here, we introduce an Expectation-Maximization (EM) algorithm to achieve a maximum likelihood estimate of the cell type composition ***μ*** by introducing *z_i,j_* as the expected probability of clustered cell type *i* coming from the real cell type *j*. Though this auxiliary variable *Z_i,j_* functions like the inverse similarity matrix *M*^-1^, its interpretation and calculation are different. Here, we could interpret *Z_i,j_* as the posterior probability of clustered cell type *i* coming from the real cell type *j*, hence can be calculated in a forward way (i.e., the E step) if given an estimated the true composition vector ***μ*** in a previous step, as follows,

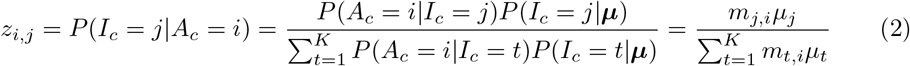

Within the EM algorithm, once we have the auxiliary variable *Z*, we can maximise the log likelihood in Eq. (1) by taking its derivative as zeros for each cell type, together with a Language multiplier on the constraint 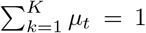. By solving these equations (i.e., M-step), we can obtain an updated ***μ*** with *Z* calculated from a previous step.

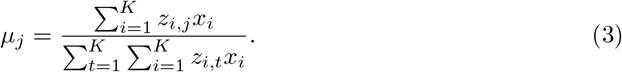

Therefore, by alternating the E step in Eq. (2) and M step in Eq. (3), we can achieve a maximum likelihood estimate of corrected cell type composition vector ***μ*** once the log likelihood stops increasing.

### 4.2 Estimation of clustering similarity matrix

As mentioned above, the clustering of single-cell is often inaccurate with systematic bias. So we introduce a similarity matrix to describe this kind of misclassification bias. We assume that this kind of bias comes from the similarity between different cell types and variety within each cell type. Thus, the conditions of samples where our single-cell data comes from do not influence the direction of this bias. In other words, misclassification error introduced by the clustering process is consistent across all samples coming from different conditions.

Currently, DCATS supports three strategies for estimating the confusion matrix: Uniform, KNN and classification. Firstly, the uniform confusion matrix is defined as

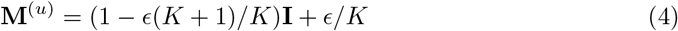

where *K* is the number of cell types; **I** is an identify matrix and *ϵ* is the level of confusion, which are set to 0. 05 by default. It describes an unbiased misclassification across all the cell types. It means that one cell from a cell type have equal chance to be assigned to rest other cell types.

Secondly, the KNN-based similarity matrix is estimated from the knn graph based on the transcriptome, e.g., provided by Seurat [18]. It calculates the proportion of neighborhoods that are regarded as other cell types. In this case, DCATS corrects cell proportions mainly based on the information of similarity between different cell types and variety within each cell types. By defining *n_x,j_* as the number of neighbors for cell *x* that are classified as cell type *j*, we can calculate KNN-based similarity matrix, e.g., its entry *m_i,j_* for similarity (misclassification) from cluster *i* to *j* as follows:

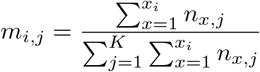

We realize in some situations, the cell clustering process might not be performed using the Seurat pipeline. It is also a common situation where people perform the clustering process based on part of the markers which they are interested in. With the development of technology, more information can be used for single cell clustering including spatial information as well as biophysical feature. In above cases, knn matrix given by the Seurat pipeline cannot fully capture the bias introduced through the clustering process.

Based on the above reason, we introduce the third way to estimate a similarity matrix. The input for estimating the similarity matrix will be a data frame containing a set of variables that the user believe will influence the result of the clustering process as well as the cell type labels for each cell. We then use 5-fold cross validation and use support vector machine as the classifier to estimate the similarity matrix.

### 4.3 Detecting differential composition abundance

#### Generalised linear model

We assume the corrected cell counts *z_s,j_* for cell type *j* given sample *s* follow beta-binomial distribution. We then describe *z_s,j_* with a beta-binomial generalised linear model(GLM) with a logit link function for each covariate *i*, as follows,

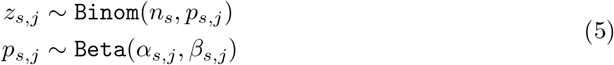

where we reparameterized with the mean *μ_s,j_* and over dispersion *ϕ_j_* as *α_s,j_* = *μ_s,j_*(1/*ϕ_j_* – 1) and *β_s,j_* = (1 – *μ_s,j_*)(1/*ϕ_j_* – 1). As described below, the mean parameter *μ_s,j_* is regressed to a set of covariates under different hypothesis. The weights for covariates and *ϕ_j_* will be optimised to achieve maximum likelihood, through the *aod* package. The p value can be calculated with a likelihood ratio test by comparing the log likelihoods in both alternative and null hypotheses.

#### Full mode and Null mode

In DCATS, we allow using both ‘full mode’ and ‘null mode’ to finish the differential abundance analysis. When using null mode, we compare these two models:

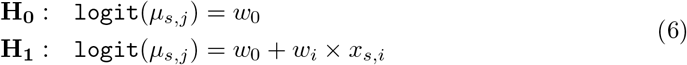

In this case, we only consider the influence of the factor we are evaluating. When using full model, we compare these two models as follows,

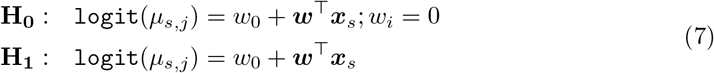

where ***x**_s_* and ***w*** are respecitvely for the weight and covariate vectors for sample *s*. Under the null hypothesis, the *w_i_* will assumed to be 0. In this case, we test the differential abundance of different condition controlling the influence of other confounding covariates.

#### Fixed overdispersion term

We found out that when the numbers of biological replicates are low, the over-dispersion term *ϕ* can be over-estimated. Thus, DCATS allows estimating a global over-dispersion term across all cell types, where we assume that the over-dispersion *ϕ_j_* remains the same across for any cell type *j*. To estimate a global dispersion term, we fit all cell types in one joint beta-binomial GLM by taking the design matrix as the cell type indicators. Then, the dispersion term estimated here is treated as a global dispersion and will be taken as a prefixed value when performing the main round GLM for each cell type.

Except for the mode we used in main text (Using KNN matrix as the similarity matrix and fixing the overdispersion term), three other modes of DCATS are tested in the simulation data (Supp. Fig. S2, Supp. Fig. S4, Supp. Fig.7-11).

- ‘wtoPhi_wtoEM’ indicates using basic beta-binomial distribution without bias correction or fixing over-dispersion term.
- ‘wtoPhi_emK’ indicates without fixing the over dispersion term Phi but clustering bias corrected by KNN matrix.
- ‘wtoPhi_emU’ indicates without fixing the over dispersion term Phi but clustering bias corrected by uniform matrix.
- ‘estPhi_wtoEM’ indicates using fixing the over dispersion term Phi but no clustering bias correction.
- ‘estPhi_emK’ indicates using fixing the over dispersion term Phi and clustering bias corrected by KNN matrix.
- ‘estPhi_emU’ indicates using fixing the over dispersion term Phi and clustering bias corrected by uniform matrix.

### 4.4 Theoretical simulation

The input of DCATS is combined with two parts. The cell count matrix *C_i_* denotes numbers of cells we observe for each cell type in each sample coming from same condition *i*. For each analysis, DCATS can only compare cell count matrices coming from two condition. Another part is the similarity matrix *M* we describe above.

We first simulate two count matrices *C*_1_, *C*_2_ directly to demonstrate that introducing the similarity matrix to correct ***μ*** is important for the differential composition analysis. We simulate two cell counts vectors ***μ***_1*s*_ for condition 1, and three cell counts vectors ***μ***_2*s*_ for condition 2. The proportions of different cell type follow a dirichlet distribution, and misclassification error is introduced based on the similarity matrix we design using multinomial distribution.

We first used DCATS and Fisher’s exact test to perform the differential composition analysis. We then used the similarity matrix we designed to correct the miscalssification error we introduced. In this case, we used the true similarity matrix to do this bias correction process. For each simulation, we repeated the above process 50 times and calculate mcc, sensitivity and specificity. Overall, we got 50 Matthews correlation coefficient (MCC), sensitivity and specificity for 50 simulation.

### 4.5 Simulation with single-cell RNA gene count matrices

When analyzing single cell RNA sequencing (scRNA-seq) data, we always start from gene expression matrix. Here we design a simulation process to mimic the process from getting gene expression matrix to analyzing differential compositions. We first using splatter [22] to generate a large cell pool including different cell types. We then generate a proportion vector 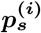 for each sample *s* from condition *i* following a Dirichlet distribution. Given a total cell count generate from a uniform distribution for each sample, a cell count vector 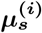 is generated from multinomial distribution.

Based on the cell count vector 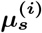, we randomly select cells as well as their gene expression profiles. We can therefore get a gene expression matrix for each sample. Then, we use Seurat [18] pipeline to do the pre-processing as well as the clustering process. The cluster annotation of each cell is treated as input for the downstrain analysis. We compare DCATS with other five tests designed for differential composition analysis which are Fisher’s exact test, scDC [10], diffcyt [21], milo [8], and speckle [20] (Supp. Fig. S1).

As milo aims to detect cell stage perturbations based on k-nearest neighbor graphs but not cell type levels, we also do the differential abundance testing on the neighborhoods level by assuming all the neighborhoods are influenced by condition if that cell types have differential proportions. In this case, we only considered neighborhoods having more than 80% of cells coming from one cell type and used neighborhoods detected by milo for testing using Fisher’s exact test, speckle [20] and DCATS.

### 4.6 Adding other confounding covariates

To demonstrate that DCATS can control the influence of confounding covariates, we designed a simulation with eight cell types. Four of them have different proportions in different conditions, while four of them have same proportions in different conditions. We also added two confounding covariates, age and gender which will influence the proportions of four among eight cell types. The influence of condition, age and gender is additive and result in the final proportion of each cell types in each biological replicate. Overall, we have 10 biological replicates in each condition with different ages and genders. We simulated proportions based on the true designed proportions from a Dirichlet distribution and cell numbers based on the simulated proportions from the multinomial distribution. Based on the simulated cell counts, we simulated gene expression matrices and did the differential abundance analysis as described previously.

### 4.7 Methods for benchmark

#### Fisher’s exact test

As Fisher’s exact test doesn’t support multiple replicates, we added the cell counts numbers of all replicates in each condition and tested based on the sum of cell counts.

#### diffcyt

diffcyt is a package design for differential discovery analyses in high-dimensional cytometry data. In order to adopt it to analyze scRNA-seq data. We first create data template using random generated flowset data, then replaced it with meta-data of data we simulate. We used default for the rest setting.

#### milo

milo aims to detect cell state changes in neighborhoods level, while both our simulated data and real-world data only have ground truth in cell type level. Thus, we linked neighborhoods with cell type based on the majority cell in that neighborhoods as what milo’s authors did in the milo paper [8]. We first used FDR<0.1 as threshold to define whether one neighborhood has cell state changes, and calculated how many more neighborhoods one condition has dominant size comparing to the other condition as well as the proportion of this difference among all neighborhoods that belong to that cell type. This proportion is regarded as perturbation level. Inferring from the Kang dataset [4] which works as the negative control, we defined that one cell type shows significant proportion changes when the perturbation level is larger than 20%. Due to the less variety of simulated data, this threshold is setted to be 0 in simulated data in order to get better performance of milo.

#### speckle and scDC

For speckle and scDC, we used default setting and standard pipeline as described in the github page/tutorials of these two tools.

## Availability of data

All datasets used in the paper are previously published and can be found in NCBI GEO database: GSE96583 (Kang2017), GSE92332 (Haber2017), and GSE158055 (COVID19).

## Availability of source code

This algorithm is implemented as an R package named DCATS which is available at https://github.com/holab-hku/DCATS.

## Acknowledgements

We thank members in Ho lab and Huang lab, especially Aaron Kwok, Weizhong Zheng, Ken Yu, Junyi Chen, for fruitful discussions. This work was supported by AIR@InnoHK administered by Innovation and Technology Commission.

## Authors’ contributions

YH and JWKH conceived the ideas and designed the study. XL developed the method, implemented the R package, and performed all the experiments. CC supported the initial development of the method, collected some data, and supported evaluation. XL, YH and JWKH wrote the paper. All authors read and approved the final manuscript.

## Ethics declarations

### Ethics approval and consent to participate

Not applicable.

### Consent for publication

All authors have approved the manuscript for submission.

