## Supplemental File for "DCATS: differential composition analysis for complex single-cell experimental designs"

### 1 Supplementary Figures and Tables

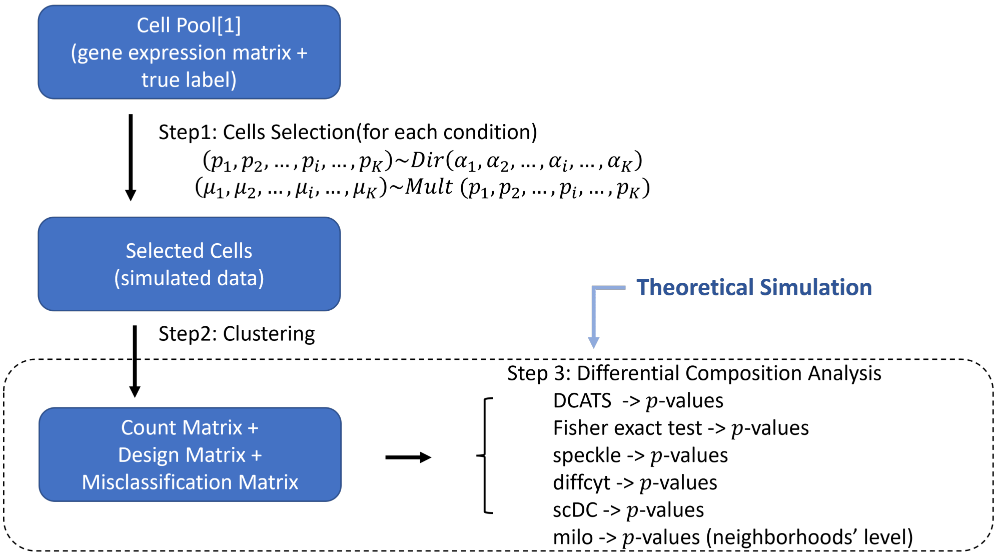


Supplementary Figure 1: Illustration of the simulation process.

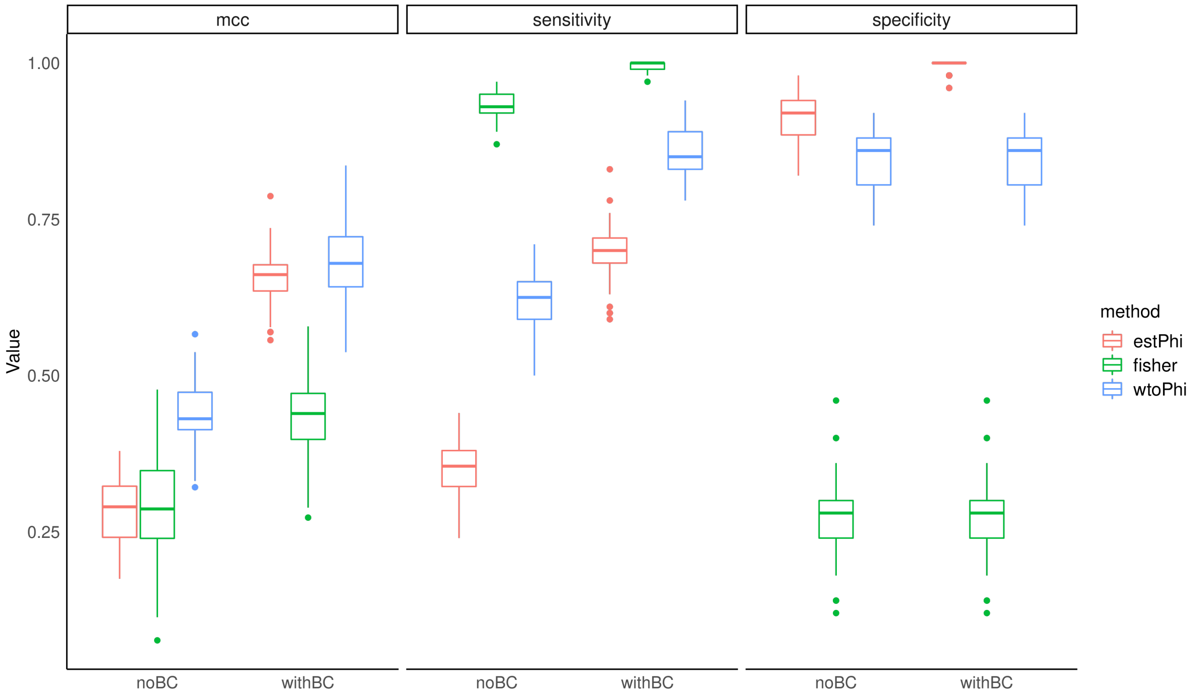


Supplementary Figure 2: The mcc, sensitivity, specificity of two DCATS models ('wtoPhi', 'estPhi') and fisher's exact test before and after bias correction.

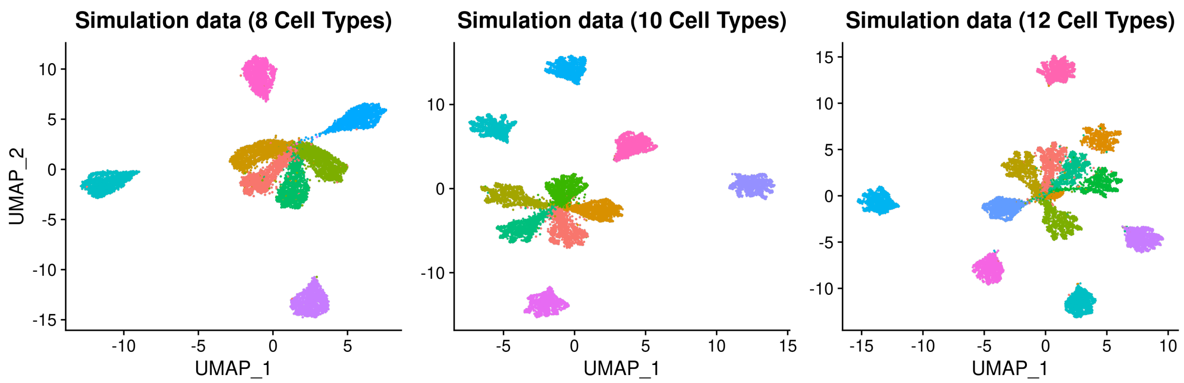


Supplementary Figure 3: The UMAP of cell pools with different numbers of cell types.

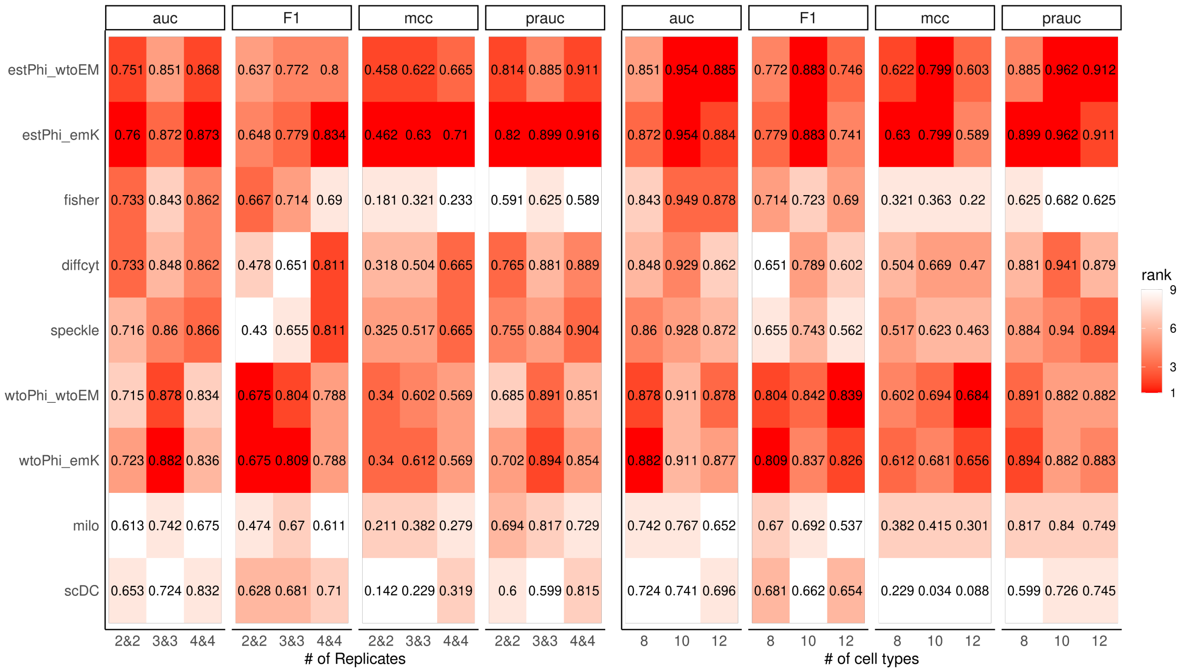


Supplementary Figure 4: The auc, F1, mcc, prauc values of different DCATS models and other methods in different simulation settings. 'wtoPhi_emK' indicates using KNN matrix to do the bias correction without pre-estimated over-dispersion term. 'wtoPhi_wtoEM' indicates using basic beta-binomial distribution without bias correction and pre-estimated over-dispersion term. 'estPhi_emK' indicates using KNN matrix to do the bias correction with pre-estimated over-dispersion term. 'estPhi_wtoEM' indicates using pre-estimated over-dispersion term without bias correction.

| | A | B | C | D | E | F | G | H |
|---|---|---|---|---|---|---|---|---|
| 7 | 0.9925884 | 0.0000000 | 0.0074116 | 0.0000000 | 0.0000000 | 0.0000000 | 0.0000000 | 0.0000000 |
| 8 | 0.0000000 | 0.9996793 | 0.0003207 | 0.0000000 | 0.0000000 | 0.0000000 | 0.0000000 | 0.0000000 |
| 3 | 0.0000000 | 0.0000000 | 0.9616307 | 0.0004796 | 0.0163070 | 0.0100719 | 0.0000000 | 0.0115108 |
| 6 | 0.0016434 | 0.0000000 | 0.0061627 | 0.9921939 | 0.0000000 | 0.0000000 | 0.0000000 | 0.0000000 |
| 2 | 0.0000000 | 0.0000000 | 0.0557157 | 0.0019212 | 0.9178674 | 0.0033622 | 0.0000000 | 0.0211335 |
| 4 | 0.0000000 | 0.0000000 | 0.0507012 | 0.0016181 | 0.0253506 | 0.9099245 | 0.0000000 | 0.0124056 |
| 5 | 0.0000000 | 0.0000000 | 0.0012804 | 0.0000000 | 0.0000000 | 0.0000000 | 0.9987196 | 0.0000000 |
| 1 | 0.0000000 | 0.0000000 | 0.1407517 | 0.0007369 | 0.0324245 | 0.0140015 | 0.0000000 | 0.8120855 |

Supplementary Figure 5: The empirical confusion matrix calculated from ground truth and seurat clustering result in one simulation with default setting.

| | A | B | C | D | E | F | G | H |
|---|---|---|---|---|---|---|---|---|
| A | 0.9985960 | 0.0000263 | 0.0010624 | 0.0000622 | 0.0001555 | 0.0000641 | 0.0000173 | 0.0000162 |
| B | 0.0000287 | 0.9991608 | 0.0006151 | 0.0000152 | 0.0001598 | 0.0000000 | 0.0000203 | 0.0000000 |
| C | 0.0018297 | 0.0009675 | 0.9375284 | 0.0021082 | 0.0187538 | 0.0094317 | 0.0006555 | 0.0287252 |
| D | 0.0000951 | 0.0000213 | 0.0018715 | 0.9972366 | 0.0001855 | 0.0003109 | 0.0000653 | 0.0002138 |
| E | 0.0003277 | 0.0003075 | 0.0229506 | 0.0002558 | 0.9404613 | 0.0119533 | 0.0000271 | 0.0237168 |
| F | 0.0001565 | 0.0000000 | 0.0133674 | 0.0004963 | 0.0138433 | 0.9546358 | 0.0000113 | 0.0174892 |
| G | 0.0000433 | 0.0000464 | 0.0009505 | 0.0001067 | 0.0000321 | 0.0000116 | 0.9987777 | 0.0000316 |
| H | 0.0000565 | 0.0000000 | 0.0582963 | 0.0004888 | 0.0393305 | 0.0250432 | 0.0000443 | 0.8767404 |

Supplementary Figure 6: The estimated knn similarity matrix calculated from ground truth and seurat clustering result in one simulation with default setting.

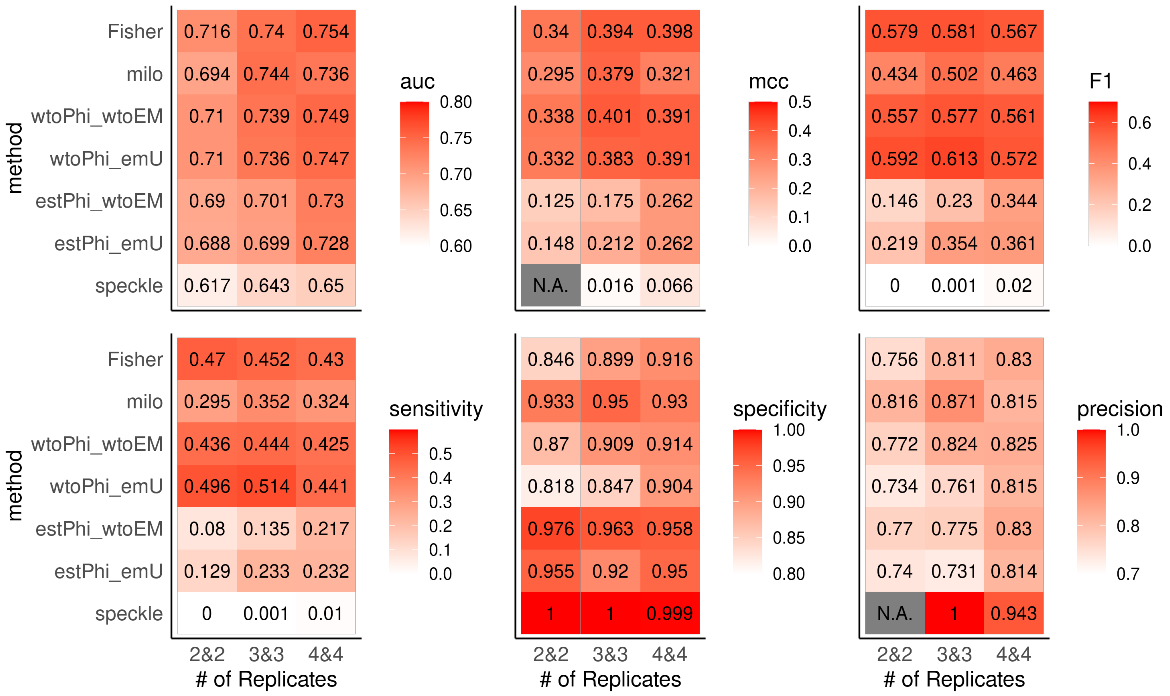


Supplementary Figure 7: The auc, mcc, F1, sensitivity, specificity and precision values of different DCATS models and other methods in simulation with different numbers of biological replicates. 'wtoPhi_emU' indicates using uniform matrix to do the bias correction without pre-estimated over-dispersion term. 'wtoPhi_wtoEM' indicates using basic beta-binomial distribution without bias correction and pre-estimated over-dispersion term. 'estPhi_emU' indicates using uniform matrix to do the bias correction with pre-estimated over-dispersion term. 'estPhi_wtoEM' indicates using pre-estimated over-dispersion term without bias correction.

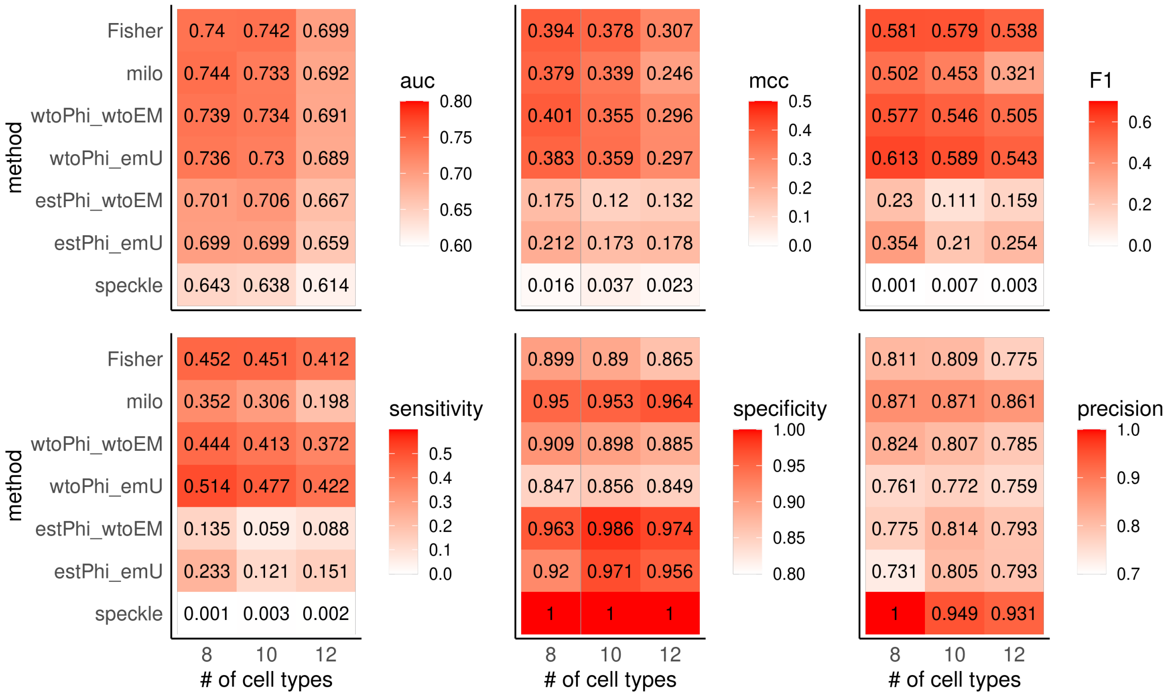


Supplementary Figure 8: The auc, mcc, F1, sensitivity, specificity and precision values of different DCATS models and other methods in simulation with different numbers of cell types. As speckle didn't detect any neighborhood which has significant proportion difference with 12 cell types scenario, the mcc and precision is 'N.A.'.

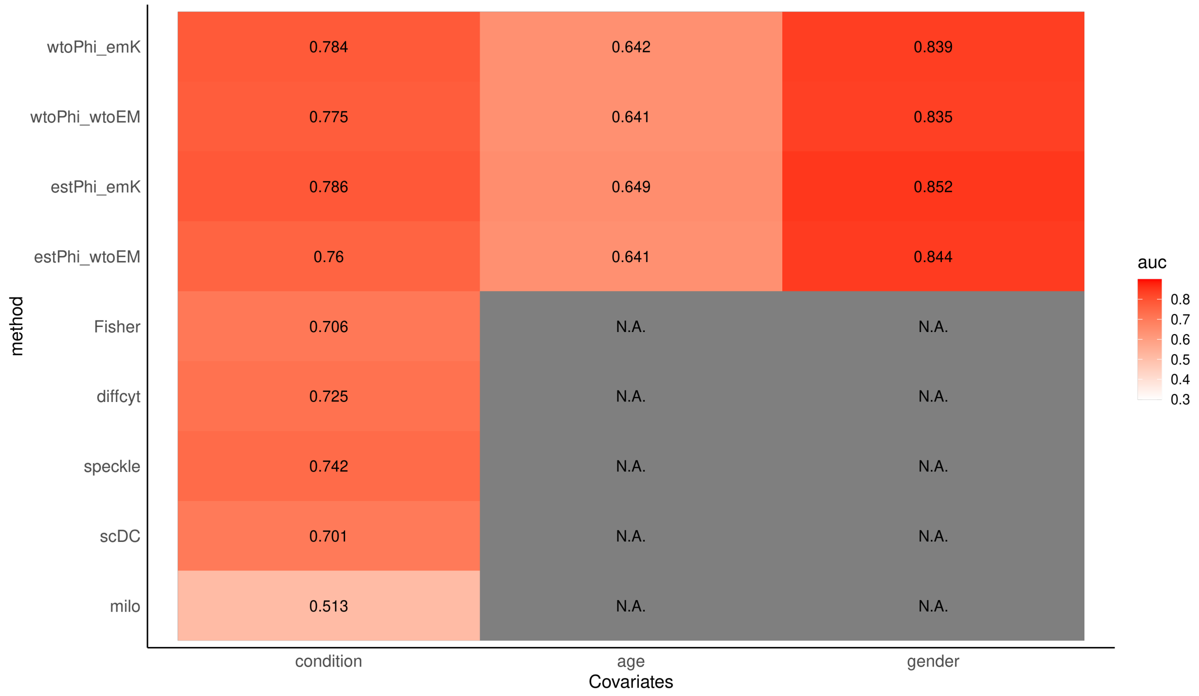


Supplementary Figure 9: The auc values of different DCATS models and other methods in the simulation with confounding covariates. 'N.A.' means not applicable (same in following two plots).

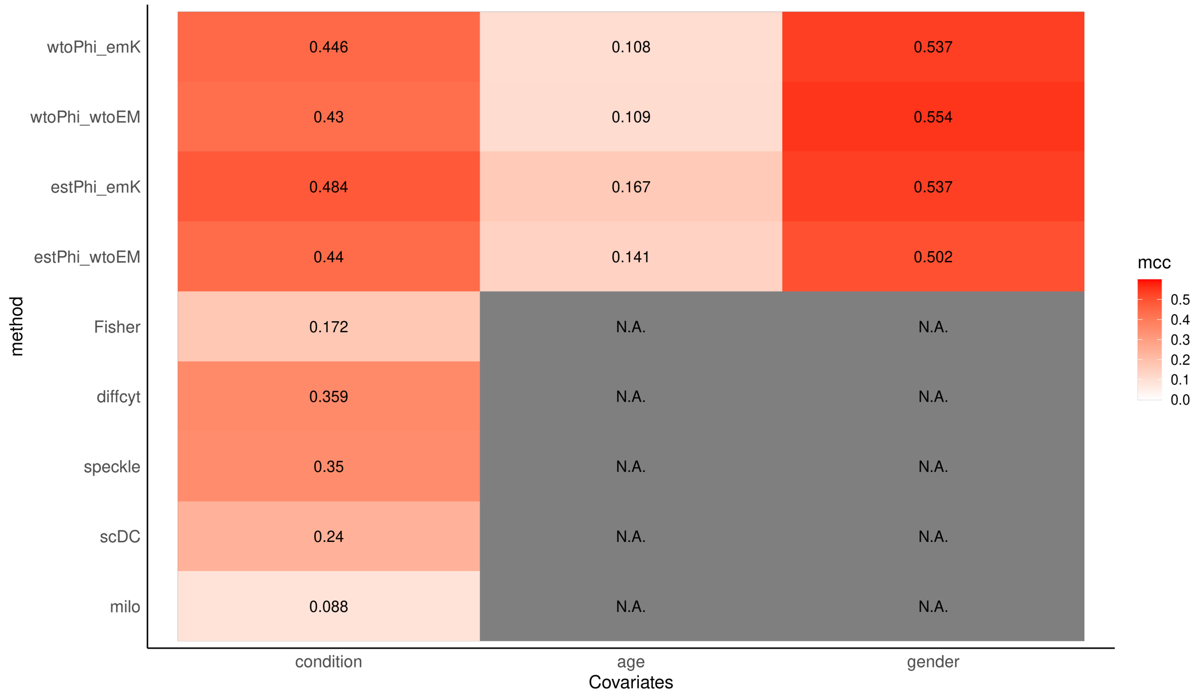


Supplementary Figure 10: The mcc values of different DCATS models and other methods in the simulation with confounding covariates.

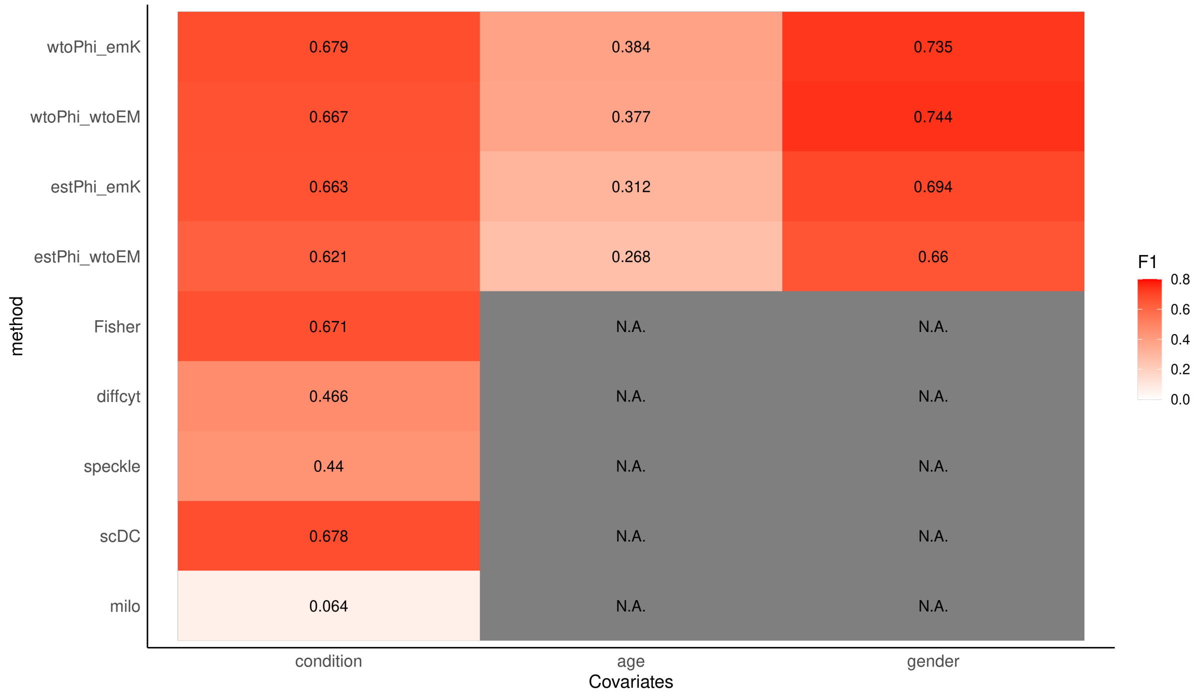


Supplementary Figure 11: The F1 values of different DCATS models and other methods in the simulation with confounding covariates.

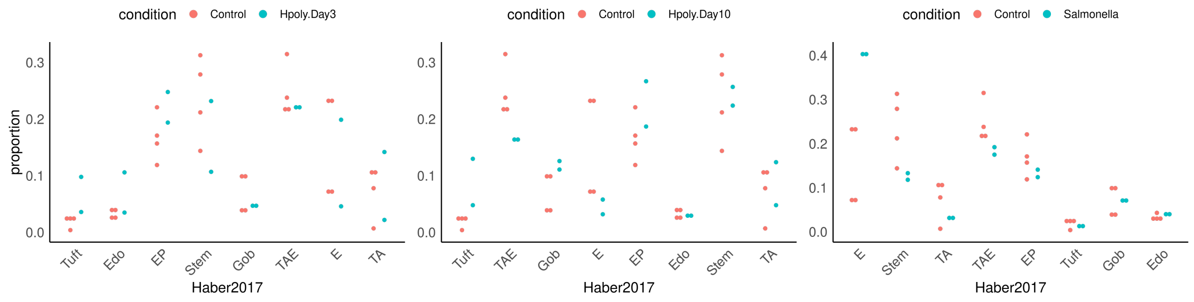


Supplementary Figure 12: The corrected cell type proportions of the Haber dataset obtained from using DCATS for bias correction

| method | mcc | auc | sensitivity | specificity | precision | F1 | replicates |
|---|---|---|---|---|---|---|---|
| estPhi_emK | **0.462** | **0.76** | 0.527 | 0.902 | 0.843 | 0.648 | 2&2 |
| estPhi_wtoEM | 0.458 | 0.751 | 0.509 | 0.911 | 0.851 | 0.637 | 2&2 |
| diffcyt | 0.318 | 0.733 | 0.339 | 0.92 | 0.809 | 0.478 | 2&2 |
| Fisher | 0.181 | 0.733 | **0.848** | 0.304 | 0.549 | 0.667 | 2&2 |
| wtoPhi_emK | 0.34 | 0.723 | 0.688 | 0.652 | 0.664 | 0.675 | 2&2 |
| speckle | 0.325 | 0.716 | 0.286 | **0.955** | **0.865** | 0.43 | 2&2 |
| wtoPhi_wtoEM | 0.34 | 0.715 | 0.688 | 0.652 | 0.664 | **0.675** | 2&2 |
| scDC | 0.142 | 0.653 | 0.732 | 0.402 | 0.55 | 0.628 | 2&2 |
| milo | 0.211 | 0.613 | 0.366 | 0.821 | 0.672 | 0.474 | 2&2 |
| wtoPhi_emK | **0.612** | **0.882** | 0.824 | 0.787 | 0.795 | **0.809** | 3&3 |
| wtoPhi_wtoEM | 0.602 | 0.878 | 0.815 | 0.787 | 0.793 | 0.804 | 3&3 |
| estPhi_emK | 0.63 | 0.872 | 0.685 | 0.926 | 0.902 | 0.779 | 3&3 |
| speckle | 0.517 | 0.86 | 0.509 | **0.954** | **0.917** | 0.655 | 3&3 |
| estPhi_wtoEM | **0.622** | 0.851 | 0.676 | 0.926 | 0.901 | 0.772 | 3&3 |
| diffcyt | 0.504 | 0.848 | 0.509 | 0.944 | 0.902 | 0.651 | 3&3 |
| Fisher | 0.321 | 0.843 | **0.972** | 0.25 | 0.565 | 0.714 | 3&3 |
| milo | 0.382 | 0.742 | 0.63 | 0.75 | 0.716 | 0.67 | 3&3 |
| scDC | 0.229 | 0.724 | 0.861 | 0.333 | 0.564 | 0.681 | 3&3 |
| estPhi_emK | **0.71** | **0.873** | 0.759 | **0.94** | **0.926** | **0.834** | 4&4 |
| estPhi_wtoEM | 0.665 | 0.868 | 0.707 | **0.94** | 0.921 | 0.8 | 4&4 |
| speckle | 0.665 | 0.866 | 0.741 | 0.914 | 0.896 | 0.811 | 4&4 |
| diffcyt | 0.665 | 0.862 | 0.741 | 0.914 | 0.896 | 0.811 | 4&4 |
| Fisher | 0.233 | 0.862 | **0.922** | 0.25 | 0.552 | 0.69 | 4&4 |
| wtoPhi_emK | 0.569 | 0.836 | 0.802 | 0.767 | 0.775 | 0.788 | 4&4 |
| wtoPhi_wtoEM | 0.569 | 0.834 | 0.802 | 0.767 | 0.775 | 0.788 | 4&4 |
| scDC | 0.319 | 0.832 | 0.888 | 0.388 | 0.592 | 0.71 | 4&4 |
| milo | 0.279 | 0.675 | 0.569 | 0.707 | 0.66 | 0.611 | 4&4 |

Supplementary Table 1: mcc, auc, sensitivity, specificity and F1 for different tests and DCATS with the change of replicates numbers. DCATS_wMSVM means using the confusion matrix calculated by support vector machine, DCATS_wMU means using the uniform confusion matrix, DCATSmeans using the confusion matrix derived from the KNN graph and DCATS_noBC means using DCATS without the bias correction step (keep onely significant decimal digits). 'replicates' indicates the number of biological replicates in each condition

| method | mcc | auc | sensitivity | specificity | precision | F1 | clustersN |
|---|---|---|---|---|---|---|---|
| estPhi_wtoEM | **0.709** | **0.91** | 0.708 | **0.975** | **0.966** | 0.817 | 8 |
| estPhi_emK | 0.703 | 0.902 | 0.725 | 0.958 | 0.946 | **0.821** | 8 |
| Fisher | 0.36 | 0.898 | **0.95** | 0.333 | 0.588 | 0.726 | 8 |
| speckle | 0.671 | 0.895 | 0.675 | 0.967 | 0.953 | 0.79 | 8 |
| diffcyt | 0.671 | 0.894 | 0.733 | 0.925 | 0.907 | 0.811 | 8 |
| wtoPhi_emK | 0.562 | 0.861 | 0.833 | 0.725 | 0.752 | 0.791 | 8 |
| wtoPhi_wtoEM | 0.562 | 0.86 | 0.833 | 0.725 | 0.752 | 0.791 | 8 |
| scDC | 0.405 | 0.807 | **0.95** | 0.383 | 0.606 | 0.74 | 8 |
| milo | 0.385 | 0.737 | 0.642 | 0.742 | 0.713 | 0.675 | 8 |
| estPhi_wtoEM | **0.697** | **0.923** | 0.707 | **0.967** | **0.955** | **0.812** | 10 |
| estPhi_emK | 0.681 | 0.92 | 0.707 | 0.953 | 0.938 | 0.806 | 10 |
| Fisher | 0.365 | 0.907 | **0.987** | 0.267 | 0.574 | 0.725 | 10 |
| speckle | 0.533 | 0.892 | 0.507 | **0.967** | 0.938 | 0.658 | 10 |
| diffcyt | 0.529 | 0.892 | 0.513 | 0.96 | 0.928 | 0.661 | 10 |
| wtoPhi_emK | 0.547 | 0.852 | 0.8 | 0.747 | 0.759 | 0.779 | 10 |
| wtoPhi_wtoEM | 0.547 | 0.851 | 0.8 | 0.747 | 0.759 | 0.779 | 10 |
| scDC | 0 | 0.698 | 0.96 | 0.04 | 0.5 | 0.658 | 10 |
| milo | 0.293 | 0.618 | 0.353 | 0.893 | 0.768 | 0.484 | 10 |
| estPhi_wtoEM | 0.589 | **0.877** | 0.667 | 0.906 | 0.876 | 0.757 | 12 |
| estPhi_emK | **0.593** | 0.874 | 0.678 | 0.9 | 0.871 | 0.762 | 12 |
| Fisher | 0.322 | 0.87 | **0.961** | 0.272 | 0.569 | 0.715 | 12 |
| speckle | 0.479 | 0.853 | 0.489 | **0.939** | **0.889** | 0.631 | 12 |
| wtoPhi_wtoEM | 0.583 | 0.844 | 0.794 | 0.789 | 0.79 | **0.792** | 12 |
| wtoPhi_emK | 0.578 | 0.843 | 0.794 | 0.783 | 0.786 | 0.79 | 12 |
| diffcyt | 0.478 | 0.836 | 0.506 | 0.928 | 0.875 | 0.641 | 12 |
| scDC | 0.112 | 0.708 | 0.85 | 0.239 | 0.528 | 0.651 | 12 |
| milo | 0.298 | 0.61 | 0.278 | 0.944 | 0.833 | 0.417 | 12 |

Supplementary Table 2: mcc, auc, sensitivity, specificity and F1 for different tests and DCATS with different numbers of cell types in samples (keep onely significant decimal digits). 'clustersN' indicates the number of cell types in samples. N.B. the actual values about 'estPhi_emK' and 'estPhi_wtoEM' are close to each other but not the same.

| cluster | truth | wtoPhi_wtoEM | wtoPhi_emSVM | estPhi_wtoEM | estPhi_emSVM | fisher | scDC | speckle | milo_pct |
|---|---|---|---|---|---|---|---|---|---|
| B cells | N | 0.997 | 0.997 | 0.995 | 0.995 | 0.338 | 0 | 0.989 | 0.032 |
| CD14+ Monocytes | N | 0.51 | 0.51 | 0.537 | 0.527 | 0.001 | 0.48 | 0.989 | 0.186 |
| CD4 T cells | N | 0.953 | 0.953 | 0.926 | 0.921 | 0.252 | 0.408 | 0.989 | 0.058 |
| CD8 T cells | N | 0.734 | 0.734 | 0.581 | 0.566 | 0.008 | 0.348 | 0.989 | 0.187 |
| Dendritic cells | N | 0.496 | 0.496 | 0.815 | 0.96 | 0.252 | 0.333 | 0.989 | 0 |
| FCGR3A+ Monocytes | N | 0.603 | 0.603 | 0.729 | 0.335 | 0.008 | 0.077 | 0.989 | 0.101 |
| Megakaryocytes | N | 0.738 | 0.738 | 0.82 | 0.364 | 0.862 | 0.761 | 0.989 | 0.167 |
| NK cells | N | 0.244 | 0.244 | 0.514 | 0.508 | 0 | 0.018 | 0.989 | 0.071 |

Supplementary Table 3: The percentages of differential abundance neighborhoods given by milo and p-values given by other methods for Kang dataset[2].

| | Edo | E | EP | Gob | Stem | TA | TAE | Tuft |
|---|---|---|---|---|---|---|---|---|
| Edo | 0.9785933 | 0.0006868 | 0.0005495 | 0.0093209 | 0.0102934 | 0.0114537 | 0.0087549 | 0 |
| E | 0 | 0.9800824 | 0.017033 | 0 | 0 | 0 | 0 | 0 |
| EP | 0 | 0.018544 | 0.9401099 | 0 | 0.0005147 | 0.0572687 | 0.0087549 | 0 |
| Gob | 0.0091743 | 0 | 0 | 0.9826897 | 0.0056613 | 0.0017621 | 0.0019455 | 0 |
| Stem | 0.0030581 | 0.0006868 | 0 | 0.0026631 | 0.8625836 | 0.0933921 | 0.0345331 | 0 |
| TA | 0 | 0 | 0.0335165 | 0 | 0.0761709 | 0.7506608 | 0.0617704 | 0 |
| TAE | 0 | 0 | 0.0082418 | 0.0039947 | 0.0303654 | 0.0748899 | 0.8793774 | 0 |
| Tuft | 0.0091743 | 0 | 0.0005495 | 0.0013316 | 0.0144107 | 0.0105727 | 0.0048638 | 1 |

Supplementary Table 4: The confusion matrix used for the Haber dataset[1].

| cluster | truth | wtoPhi_wtoEM | wtoPhi_emSVM | estPhi_wtoEM | estPhi_emSVM | fisher | scDC | speckle | milo_pct | treatment |
|---|---|---|---|---|---|---|---|---|---|---|
| Endocrine | N | 0.13 | 0.13 | 0.288 | 0.273 | 0 | 0.36 | 0.714 | 0 | Hpoly.Day3 |
| Enterocyte | N | 0.641 | 0.641 | 0.448 | 0.41 | 0 | 0.002 | 0.748 | 0.071 | Hpoly.Day3 |
| Enterocyte.Progenitor | N | 0.103 | 0.103 | 0.315 | 0.278 | 0 | 0.341 | 0.714 | 0.067 | Hpoly.Day3 |
| Goblet | N | 0.437 | 0.437 | 0.589 | 0.571 | 0.002 | 0.003 | 0.748 | 0 | Hpoly.Day3 |
| Stem | N | 0.254 | 0.254 | 0.222 | 0.164 | 0.032 | 0.01 | 0.714 | 0 | Hpoly.Day3 |
| TA | N | 0.883 | 0.883 | 0.883 | 0.975 | 0.205 | 0.099 | 0.993 | 0 | Hpoly.Day3 |
| TA.Early | N | 0.288 | 0.288 | 0.64 | 0.612 | 0.002 | 0.006 | 0.781 | 0.088 | Hpoly.Day3 |
| Tuft | P | 0.04 | 0.04 | 0.07 | 0.061 | 0 | 0.069 | 0.714 | 0 | Hpoly.Day3 |
| Endocrine | N | 0.39 | 0.39 | 0.925 | 0.867 | 0.379 | 0.035 | 0.923 | 0 | Hpoly.Day10 |
| Enterocyte | P | 0.091 | 0.091 | 0.009 | 0.006 | 0 | 0.002 | 0.149 | 0.472 | Hpoly.Day10 |
| Enterocyte.Progenitor | N | 0.132 | 0.132 | 0.285 | 0.233 | 0 | 0.103 | 0.349 | 0.162 | Hpoly.Day10 |
| Goblet | P | 0.051 | 0.051 | 0.116 | 0.105 | 0 | 0.009 | 0.235 | 0.217 | Hpoly.Day10 |
| Stem | N | 0.824 | 0.824 | 0.865 | 0.83 | 0.279 | 0.406 | 0.923 | 0.096 | Hpoly.Day10 |
| TA | N | 0.852 | 0.852 | 0.836 | 0.592 | 1 | 0.572 | 0.923 | 0.647 | Hpoly.Day10 |
| TA.Early | P | 0.015 | 0.015 | 0.132 | 0.083 | 0 | 0.224 | 0.235 | 0.266 | Hpoly.Day10 |
| Tuft | P | 0.02 | 0.02 | 0.013 | 0.01 | 0 | 0 | 0.112 | 0.5 | Hpoly.Day10 |
| Endocrine | N | 0.426 | 0.426 | 0.798 | 0.784 | 0.429 | 0.84 | 0.761 | 0.308 | Salmonella |
| Enterocyte | P | 0.008 | 0.008 | 0 | 0 | 0 | 0.022 | 0.008 | 0.457 | Salmonella |
| Enterocyte.Progenitor | N | 0.146 | 0.146 | 0.561 | 0.495 | 0 | 0.357 | 0.647 | 0.019 | Salmonella |
| Goblet | N | 0.681 | 0.681 | 0.821 | 0.802 | 0.558 | 0.253 | 0.864 | 0.1 | Salmonella |
| Stem | P | 0.031 | 0.031 | 0.059 | 0.046 | 0 | 0.08 | 0.094 | 0.32 | Salmonella |
| TA | P | 0.026 | 0.026 | 0.192 | 0.318 | 0 | 0.801 | 0.216 | 0.538 | Salmonella |
| TA.Early | P | 0.03 | 0.03 | 0.261 | 0.268 | 0 | 0.39 | 0.216 | 0.015 | Salmonella |
| Tuft | N | 0.587 | 0.587 | 0.833 | 0.844 | 0.129 | 0.059 | 0.761 | 0 | Salmonella |

Supplementary Table 5: The percentages of differential abundance neighborhoods given by milo and p-values given by other methods for Haber dataset[1].
